# Development of a long-read high-throughput chromatin conformation capture method for evaluating both *cis* and *trans* interactions between integrated Hepatitis B Virus and host genomes

**DOI:** 10.1101/2025.10.21.683758

**Authors:** Yih-Ping Su, Joshua P. Earl, Azad A. Ahmed, Bhaswati Sen, Samuel Czerski, Garth D. Ehrlich

**Author notes:** Corresponding author: Garth D. Ehrlich, PhD, FAAAS, FAAM, FILADS 245 N. 15^th^ street, NCB 5th Floor, Philadelphia, PA 19102.

## Abstract

High-throughput chromatin conformation capture (Hi-C) has been utilized for characterizing the 3-dimensional (3D) interactome of the genome. However, short-read sequencing approaches have limited the information it provides, especially for integrated proviral DNAs . Herein we describe the development of a pipeline to investigate the 3D interactome associated with integrated Hepatitis B Virus (HBV) DNAs (iDNAs) to further understand the role of HBV iDNA in liver carcinogenesis. We employed long-read DNA sequencing combined with a novel target-pull-down library construction approach that protects against DNA shearage resulting in greatly improved sensitivity and specificity for the targeted HBV iDNA. Most importantly, we successfully increased the length of captured Hi-C library to the multi-kilobase-level and report Hi-C contacts based on a custom-developed informatics pipeline. We anticipate this combined laboratory-informatics pipeline will provide a more comprehensive view of HBV iDNA-host genomic interactions associated with their role in liver carcinogenesis, but even more importantly that it should be generalizable for the study of the geno-topological effects associated with any type of integrated DNA. (word count: 164/250)

## INTRODUCTION

DNA tumor viruses such as Human Papilloma Virus (HPV) and Hepatitis B Virus (HBV) often integrate into the host genome (Buchinsky et al. 2019; Xu et al. 2024; Ma et al. 2025) and this integration has been associated with carcinogenesis. Hepatocellular carcinoma (HCC) is one of the leading causes of cancer deaths in the United States and the world. Due to the lack of effective early detection and treatment options, HCC only has an 18% 5-year survival rate which drops to ∼ 2% following metastasis (Tran et al. 2023). More than 50% of the HCC cases reported worldwide are associated with chronic HBV infection (Parkin 2006). HBV belongs to the *Hepadnaviridae* family, genus *Orthohepadnavirus*, and has a DNA genome size of ∼ 3.2 kb (Summers et al. 1975). Although DNA integration is not required for HBV to complete its lifecycle, it has been found that integration events occur relatively early during the course of infection with a frequency of about 1 in 1,000 cells within 7 days post-infection in cell line settings (Yang and Summers 1999; Summers et al. 2003; Mason et al. 2005; Tu et al. 2018). More than 85% of the HBV-related HCC (HBV-HCC) cases harbor detectable levels of integrated HBV DNAs (HBV iDNAs). This is significantly higher than the 30-40% level observed for noncancerous HBV-related chronic hepatitis and cirrhosis cases (Koshy et al. 1981; Shafritz et al. 1981; Scotto et al. 1983; Murakami et al. 2005). Importantly, even though HBV DNA integration does not target specific regions of the host genome, it has been reported that HBV iDNA accumulates in transcriptionally active regions of the host genome and there are reports of recurrent sites adjacent to known oncogenes suggesting potential disruption of normal host gene expression patterns (Twist et al. 1981; Ding et al. 2012; Sung et al. 2012; Li et al. 2014; Zhao et al. 2016; Moreau et al. 2018; Yang et al. 2020; Tang et al. 2021; Dias et al. 2022; Guo et al. 2023; Yang et al. 2024). Expression of HBV surface antigen (HBsAg) is one of the major risk factors for liver carcinogenesis; moreover, it has been shown that HBV iDNA is the major source of HBsAg production in patients under successful antiviral treatment (Wooddell et al. 2017). Collectively, these evidences strongly suggest that HBV iDNA plays an important role in HBV-related liver carcinogenesis.

Aside from the direct interruption HBV iDNA brings to the host cell’s genome, it is unknown if integration affects the 3-dimensional (3D) topology of the genomic architecture such that novel DNA regions, in *trans*, become interacting and affect gene expression at noncontiguous distant sites. It is known that other viruses can interact with the host genome and alter the 3D structure and epigenome of the host genome thereby affecting the host cellular environment (de Bustros et al. 1988; Friedman et al. 2022). Several studies have utilized high-throughput chromatin conformation capture (Hi-C) together with short-read DNA sequencing to dissect the interactions between the HBV and host genomes (Moreau et al. 2018; Yang et al. 2020; Tang et al. 2021; Guo et al. 2023). However, these approaches are unable to differentiate among the various HBV DNA forms which in addition to HBV iDNA include relaxed-circular DNA (rcDNA), covalently closed circular DNA (cccDNA), and double-stranded linear DNA (dslDNA) which all reside in the nucleus of the HBV infected cells and share nearly identical sequences.

Since Hi-C was developed for short-read sequencing platforms that mostly provide 150 bp pair-end reads to detect potential interactions, these short-read studies can only suggest 3D *trans* interactions between the host genome and iDNA unless the evaluated environment only contained HBV iDNA and was supported by whole-genome sequencing (WGS) data confirming the structure of the HBV iDNA and its flanking genomic regions. With such complications, although HBV iDNA has been recognized as one of the major carcinogenic factors and genomic interactions between HBV DNA and host genome have been reported, it is still not well understood whether HBV iDNA is also interacting with the host genome in a way that affects the host 3-D chromatin structure. To address this question and overcome the previous technical limitations mentioned above, we developed an assay that incorporated Hi-C, HBV-targeted large-fragment DNA capture, and long-read sequencing that could directly characterize both *cis* and *trans* interactions between iDNA and the human genome. This approach provided for HBV-targeted DNA sequencing read­lengths of >> 3 kb allowing for complete *cis* (integration) and t*rans* (topological variants) characterizations with the host genome.

## RESULTS

### Restriction endonuclease substitution

To achieve the goal of adapting Hi-C technology to the PacBio long-read platform for characterizing integrated proviral DNA interactions with the host genome, it was first necessary to make modifications at the restriction enzyme (RE) digestion step to increase the length of the DNA fragments generated. This increased target sequence length provided for both complete characterization of *cis* integration sites (5’ and 3’ flanking sequences) and Hi-C-ligated products created at discontinuous topologically-interacting sites. This required the choice of a RE that did not cut within the HBV iDNA and was also a rare cutter of the human genome with a long recognition sequence. Moreover, to fit into the purchased Phase Genomics’ (PG) standard Hi-C protocol, the replacement restriction enzyme needs to have at least 1 A nucleotide in the 5’ overhang. Based upon the National Center for Biotechnology Information (NCBI)-available Reference Sequence (RefSeq) collection’s HBV genome sequence (NC_003977.2); published articles; and HBV integration related digestion protocols, *Hind*III was selected as our top candidate (Brechot et al. 1980; Edman et al. 1980; Lieberman-Aiden et al. 2009).

### In silico evaluation of Hind*III*

To compare the performance of *Hind*III and the default restriction enzyme (*Dpn*II) from the PG kit, an R package, DECIPHER, was used to computationally digest the human genome with the two enzymes (Wright 2016; Team 2020). The RefSeq of human genome (hg38) downloaded from NCBI was provided to DECIPHER as well as the HBV RefSeq genome to serve as the digestion targets. The program returned the location of detected restriction sites on each given chromosome. We then counted the total number of restriction sites and the length of the digested products. The simulated digestion results showed that HBV possesses no *Hind*III sites, while there are 9 *Dpn*II sites (**Table 1**). When evaluating the total number of fragments produced when digesting a combination of human chromosomes and HBV genome, compared with *Hind*III digestion, *Dpn*II digestion produced nearly 10 times as many fragments. The average fragment length of the digested fragments showed that *Hind*III yielded fragments nearly ten times longer on average than *Dpn*II.

**Table 1.** In silico digestion results of hbv and human genome.

| Restriction<br>Enzyme | Cuts in HBV<br>(n) | Total Fragments<br>(n) | Mean Fragment Length<br>(bp) |
| --- | --- | --- | --- |
| *DpnII* | 9 | 7,199,468 | 429 |
| *HindIII* | 0 | 860,390 | 3589 |

Overall, *in silico* digestion of the human chromosomes and HBV genome demonstrated that the *Hind*III is a non-cutter of the HBV genome, a relatively rare-cutter against human chromosomes, and most importantly, yielded large DNA fragments suitable for evaluating HBV DNA interactions at noncontiguous sites.

### Cell line selection for Hi-C SMRT assay development

Early studies using the restriction endonuclease *Hind*III, which does not cleave HBV genome, and *EcoR*I found a range of 3 to 6 HBV iDNA sequences in PLC/PRF/5 (PLC) cells (Brechot et al. 1980; Edman et al. 1980; Marion et al. 1980; Koshy et al. 1981). By using whole genome sequencing (WGS) technologies including Illumina and PacBio at least 7 different HBV iDNA copies have been documented in PLC cells (Meng et al. 2019; Chen et al. 2021; Ramirez et al. 2021; Guan et al. 2024). Therefore, we selected the PLC cell line for its well characterized HBV iDNA copies and its accessible PacBio-generated long-read WGS data. The multiple HBV iDNA copies present in PLC cells also provided an excellent target for assay development compared to other cell lines containing fewer copies of HBV iDNA. In addition to PLC cells, we also used another established liver cancer cell line, HepAD38, which was established by transfecting a plasmid containing one copy of HBV pregenomic RNA (pgRNA) cDNA sequence under the control of tetracycline-responsive promotor into HepG2 cells (Ladner et al. 1997). Upon the removal of tetracycline (tet), HepAD38 can support HBV replication. Thus, we included HepAD38 cultured cells under tet selection as our negative control, mimicking the environment where there are HBV sequences but no HBV iDNA present.

### Construction of Hi-C and Hi-C SMRT libraries

A Hi-C library constructed from PLC cells was prepared using the standard PG Hi-C protocol to serve as a method control. Since PacBio’s SMRT library construction is not optimized for on-bead library preparation, we tried heat treatment and phase separation to dissociate captured Hi-C template from streptavidin beads. However, neither of these separation methods provided sufficient yield for SMRT library construction. We then used the yield number from our standard Hi-C preparation and calculated back to estimate the yield post-Hi-C capture under standard procedure. Taking the post-capture amplification cycles into account, we confirmed the low yield was expected and that we would need a bridging step to allow SMRT library construction. To compensate for the low yield after capture and connect Hi-C construction with SMRT library construction, we worked with PacBio to incorporate the adaptor amplification processes for their long-read technology. The adaptor ligation and amplification processes involved in the incorporated protocol were very similar to Illumina’s on-bead library preparation. We ligated the amplification adaptor containing both the universal sequence for PCR amplification and a PacBio specific barcode sequence for sample pooling needed for the captured bead-bound Hi-C library. Then, a high-fidelity DNA polymerase specialized for long amplicon production (LA Taq DNA polymerase, TaKaRa, San Jose, CA) was used for amplification with an extended elongation time to ensure full-length copying and to obtain sufficient yield from the bead-bound Hi-C library templates. This adaptation successfully avoided steps for biotin-streptavidin separation and kept the overall protocol similar to the standard protocol, while most importantly increasing the final yield necessary for the minimum input for SMRT library construction. Finally, SMRTbell adaptors were added onto the Hi-C prepared DNA following the PacBio HiFi express template preparation kit 2.0 to complete the preparation of Hi-C SMRT library. The Hi-C SMRT library constructed was then analyzed on both a Qubit and an Agilent TapeStation for determination of concentration and library size distribution, respectively. By comparing the pre-sequencing quality check reports of the standard Hi-C library and the modified Hi-C SMRT library, the size of the final constructed library showed a ten-fold increase from an average length of ∼ 600 bp to > 6 kb, indicating that our protocol modifications successfully fulfilled the aim of increasing library fragment size (**Figure 1**). The constructed Hi-C SMRT library was then sequenced on a PacBio Sequel IIe with a 30-hour movie time. The run report indicated that it was highly successful with a P1 (Poisson distribution of one DNA molecule per ZMW) of 62%, 1,100,119 of HiFi reads, and an average HiFi read length of 3.8 kb. This initial Hi-C SMRT run is referred to as the OG run in the later sections when compared to other runs. Both the pre-and post-sequencing QC suggested that the modifications we made to increase the length of the Hi-C library were highly successful.

**Figure 1.**
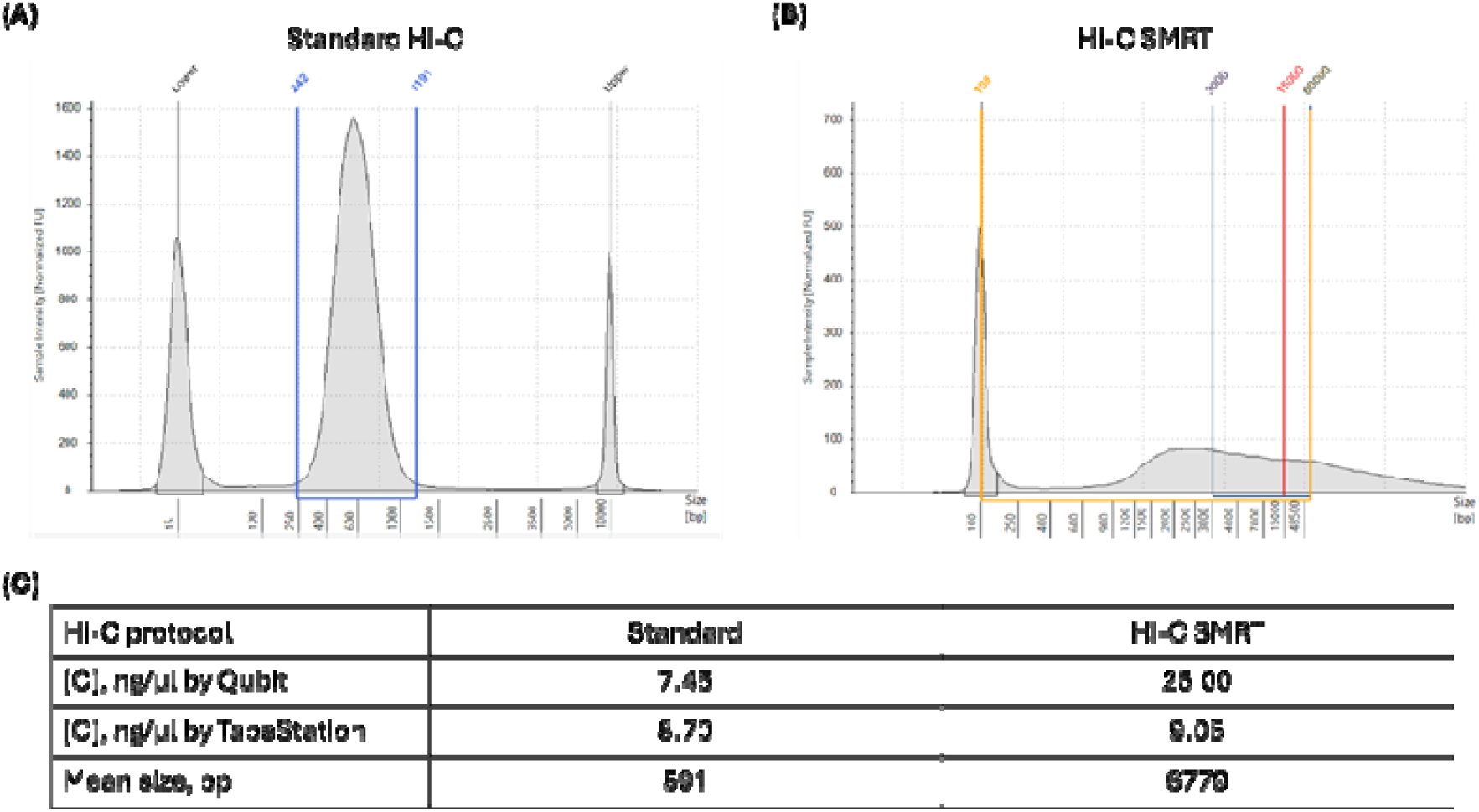
Size and concentration of the constructed Hi-C libraries. (A) Size distribution of the standard Hi-C library constructed. (B) Size distribution of the Hi-C SMRT library constructed. (C) Concentrations and mean library sizes of the constructed standard Hi-C library and Hi-C SMRT library. [C] = concentration.

### Alignment of OG run reads to reference genomes

Reads were first aligned to the genome references using minimap2. The genome references used were a combination of the human genome RefSeq (hg38) and HBV genome RefSeq (NC_003977.2) downloaded from NCBI. After alignment, the output binary alignment map (BAM) file was sorted, filtered, and indexed using samtools in preparation for downstream analyses. After we filtered for HBV sequence containing reads, we were only able to locate one HBV-containing read. With such a low sensitivity towards HBV-containing sequences we inferred that HBV-targeted capture would be essential for assay sensitivity towards HBV.

### Target-enrichment for long DNAs

Traditional hybridization capture methods for the production of pull-down libraries were designed for short-read sequencing platforms. However, these capture strategies are not suitable for long DNA fragments as they are prone to shearing during the traditional hybridization capture process and are thus not suitable for Hi-C SMRT libraries that are composed of long DNA fragments. To ensure coverage for all major HBV genotypes we submitted 42 HBV reference genomes (to provide coverage for the 10 classified major genotypes of HBV (A – J) and their variants) to Twist Bioscience (Carlsbad, CA) for the design and construction of a comprehensive set of HBV hybridization probes (**Supplementary Table 1** and **Figure 2**) (McNaughton et al. 2020). This probe set was purchased together with Twist Bioscience’s hybridization capture kit that was specifically designed for the long-read PacBio platform. To assess the effectiveness of the incorporated HBV-targeted capture, we prepared freshly harvested PLC cells and split them into two equal aliquots. Both aliquots went through Hi-C construction, adaptor ligation, and amplification. Then one of the libraries was subjected to HBV-targeted capture followed by SMRT library construction (UA2) while the other aliquot went straight to SMRT library construction (UA1). The libraries were evaluated as above with both having much larger average size distributions than the 5 kb expected (**Figure 3**). These two individually bar-coded libraries were then sequenced together as a pool.

**Figure 2.**
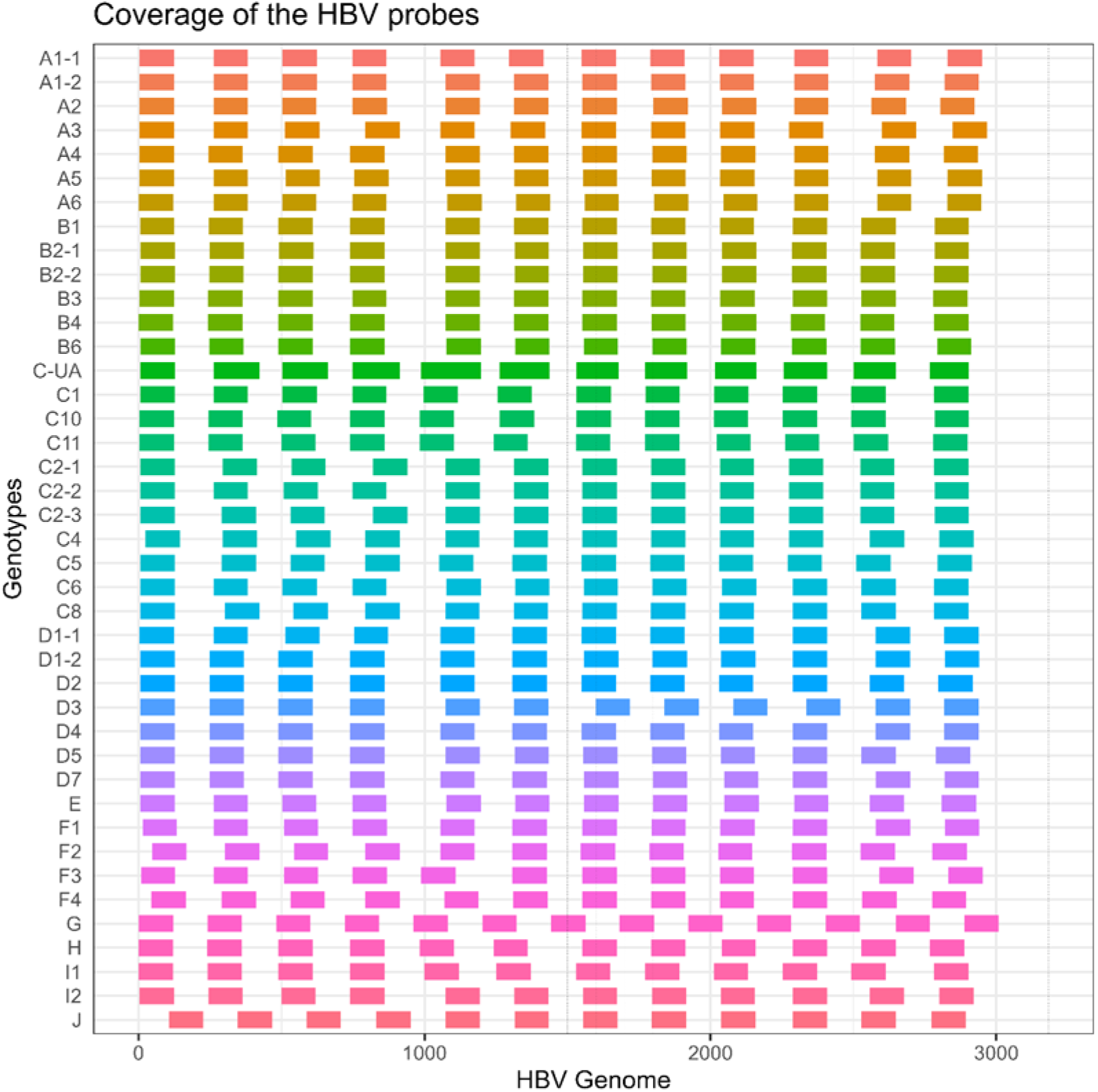
HBV hybridization probe map. Probes were designed by Twist Bioscience based on the 42 HBV genotypes suggested by McNaughton et al. 2020 which had been visualized using R package ggplot. Each row represents one HBV genotype in the same order as that of Supplementary Table 1. Each probe is illustrated as solid line colored by genotype, showing the location of the probes and the regions covered.

**Figure 3.**
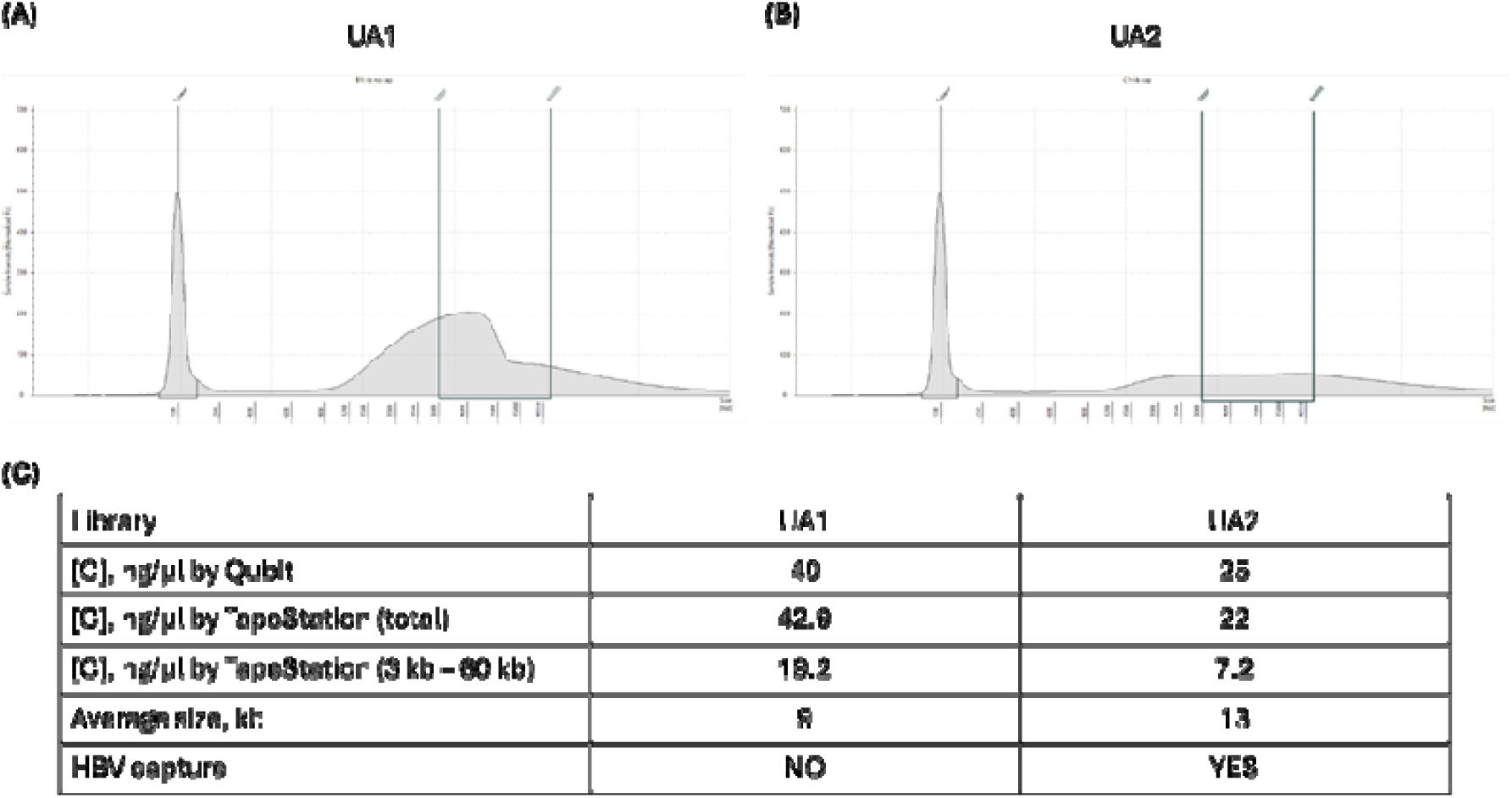
Size and concentration of the constructed Hi-C SMRT libraries. (A) Size distribution of the constructed Hi-C SMRT library, UA1. (B) Size distribution of the constructed Hi-C SMRT library, UA2. (C) Average library size and concentrations of the constructed Hi-C SMRT libraries. [C] = concentration.

### Analyzing the HBV-Hi-C SMRT data

The pooled library gave an excellent P1 of 76.8%, indicating a very successful sequencing run. As expected, there were a greater number of reads from the non-captured library as the pool itself was skewed due to the limited DNA available from the captured library (**Figure 4**). However, when we examined the read content, the captured library contained > 50% of HBV-containing reads which is much higher than that of the non-captured library (>0.001%) (**Figure 4**). We assessed the length of the aligned reads from the OG, UA1, and UA2 libraries, demonstrating that the 3 Hi-C SMRT libraries all had major length populations in the 3kb rang (**Table 2**). Overall, the data from the three libraries demonstrated high levels of consistency among the three Hi-C SMRT preparations, but most importantly showed the efficiency of incorporating an HBV-targeted capture step.

**Figure 4.**
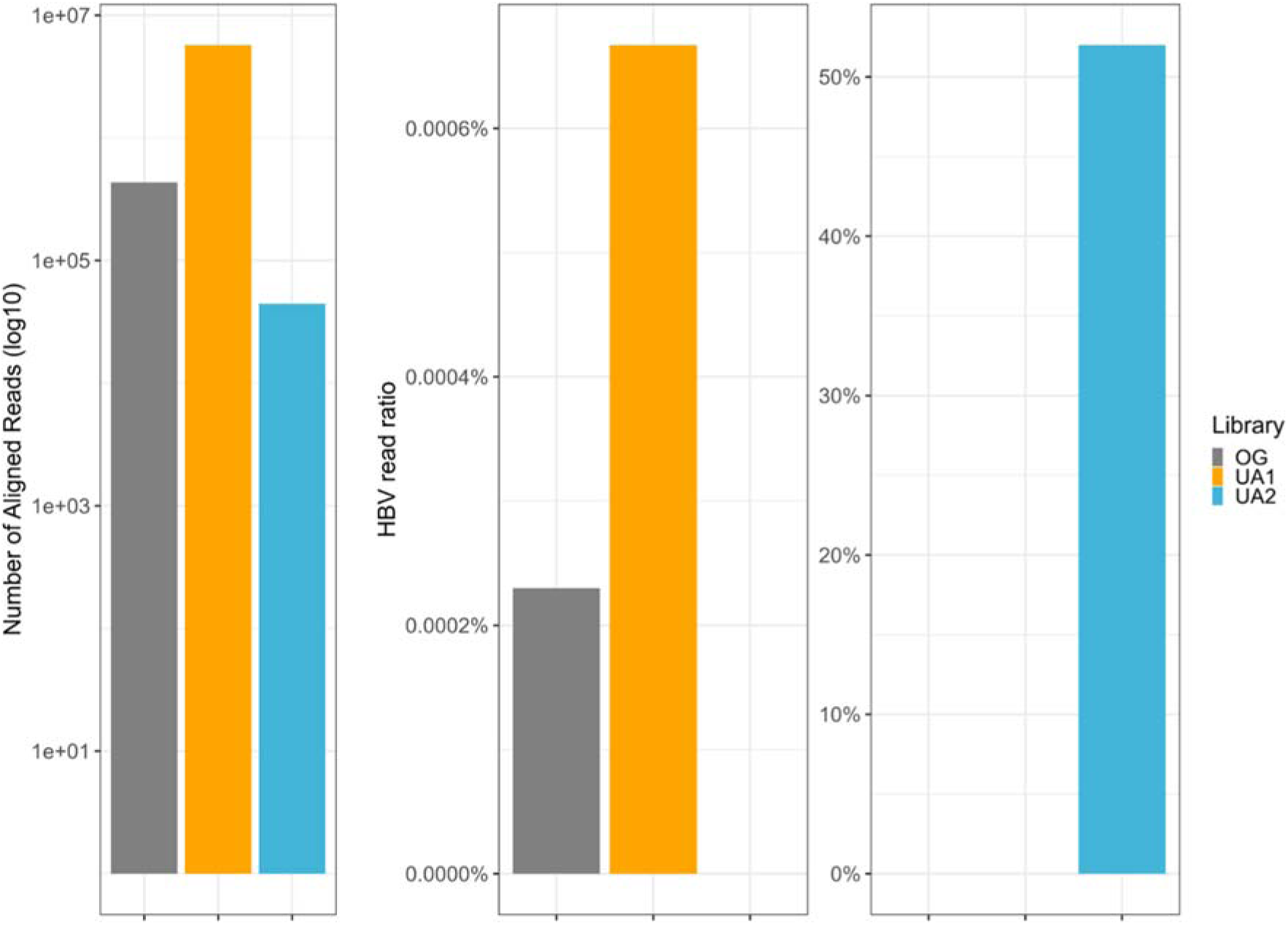
Reads of the constructed Hi-C SMRT libraries. The leftmost panel shows the number of reads aligned to a combined reference of HBV and human genome from each Hi-C SMRT library on a log scale. The middle panel shows the percentage of reads aligned to HBV from library OG and UA1, which both are below 1%. The panel on the right shows the percentage of HBV-aligned reads from UA2.

### Findings in HepAD38 Hi-C SMRT library

As described in the establishment of cell line HepAD38, it was developed by transfecting an HBV-free liver cancer cell line with plasmid, ptetHBV, which was constructed on a pBR322 backbone (Aden et al. 1979). The sequence information of the backbone pBR322 was obtained (NEB, Ipswich, MA) and only one *Hind*III restriction site on the backbone was identified. The single restriction site located on the plasmid backbone made it a good control for non-integrated HBV DNA. Since the *Hind*III does not cut the HBV genome, thus the HBV Hi-C reads identified in HepAD38 should also contain traces of pBR322 adjacent to the *Hind*III restriction site. Thus, we harvested HepAD38 and performed the same Hi-C SMRT process including adaptor ligation amplification, HBV-target capture, and sequencing of the constructed library as described in the previous sections. Encouragingly, a total number of 781,744 reads from this preparation were obtained. A total of 258,509 reads were aligned to either or both the HBV and human genome RefSeq and 1,215 of them were HBV-containing reads (**Table 2**). At first glance, 1,215 HBV-containing reads seem to be few for an HBV-target captured library. However, the ratio of HBV-containing reads, calculated by dividing the number of HBV-containing reads by the number of aligned reads, showed that the HBV-target capture worked well and reached 0.47% compared to those non-captured libraries with ratios below 0.01%. To check if potential Hi-C reads were obtained, we also searched for duplicated *Hind*III restriction sites (dRE), which are characteristic of the Hi-C ligation process, within the HBV-containing reads. Although the number of HBV-containing reads harboring dRE sequences were low in HepAD38 library, the percentage of dRE detected in HBV-containing reads suggested that the HBV-target capture not only increased the assay sensitivity towards HBV specific sequences, but also consequently increased the sensitivity towards potential HBV Hi-C reads. Overall, the sequencing results of all Hi-C SMRT libraries showed consistent success in delivering longer read lengths compared to standard Hi-C protocols. Moreover, both libraries (UA2, HepAD38) that were HBV-target captured demonstrated the importance and success of target enrichment with higher percentages of HBV-containing reads and dRE-HBV-containing reads (**Table 2**).

**Table 2.** Read number breakdown of the hi-c smrt libraries.

| Library | Cell line | Mean RL<br>(bp) | HBV<br>(n) | HBV<br>(%) | dRE<br>(n) | dRE<br>(%) |
| --- | --- | --- | --- | --- | --- | --- |
| OG | PLC | 2,337 | 1 | 0.000 | 0 | 0 |
| UA1 | PLC | 4,104 | 38 | 0.001 | 0 | 0 |
| UA2 | PLC | 4,002 | 23,036 | 51.987 | 2,900 | 12.59 |
| HepAD38 | HepAD38 | 3,754 | 1,215 | 0.470 | 70 | 5.76 |

Legend: RL = read length; dRE = double restriction sites.

### Tools to identify potential Hi-C SMRT reads in PLC cell line

Following the development of the technical protocols to produce the Hi-C SMRT sequence libraries, we then developed a bioinformatic pipeline to dissect and analyze the performance of each Hi-C SMRT run. Since all of our initial Hi-C SMRT libraries were constructed using PLC cells, we downloaded the PacBio-produced whole genome sequencing (WGS) files for PLC from the Sequence Read Archive (SRA) at NCBI to serve as a reference (Chen et al. 2021). Due to the fact that nearly all of the existing Hi-C analysis tools were developed for standard short-read sequencing protocols, we explored alternative tools and methods to evaluate our long-read Hi-C SMRT libraries. Although these short-read methods were not ideal for analyzing long sequencing data such as our Hi-C SMRT data, these standard Hi-C analytic tools served as references for the principles and processes needed to analyze the long-read Hi-C SMRT data. Our initial evaluations showed that converting Hi-C protocols from short-read sequencing platforms to long-read sequencing platforms resulted in sacrificed coverage levels. However, through optimization of both technical and informatic protocols we have produced 3D contact data and anticipate that future verifications experiments will support our findings.

### Read compositions of the datasets

Sequence alignment was performed using minimap2 with a custom-prepared merged reference containing both the human genome RefSeq (hg38) and the HBV genome RefSeq (NC_003977.2) where the HBV genome was added as an additional chromosome to the human genome. The full alignment output was saved as a BAM file containing information including read names, flags (mapping result), reference name, reference coordinates, concise idiosyncratic gapped alignment report (CIGAR), and mapped sequences. From the BAM file, we filtered out those reads that were not mapped to the references and extracted the mapped reads. Likewise, further filtering was performed to extract the subset of reads with sequences mapping to HBV. With these subset read files, we were able to count and summarize the number of each type of read in each library (**Table 3**). The percentage of HBV-containing reads was calculated by dividing the number of mapped HBV-containing reads by the total number of mapped reads. The number of reads containing double restriction sites (dRE), indicative of Hi-C ligations, was counted by filtering reads with dRE specific sequences. Notably, the filtering process for identification of the dRE pattern was performed by simple text search due to the short length of the restriction site (< 10bp) which was too short to serve as reference sequence for standard alignment programs.

**Table 3.**
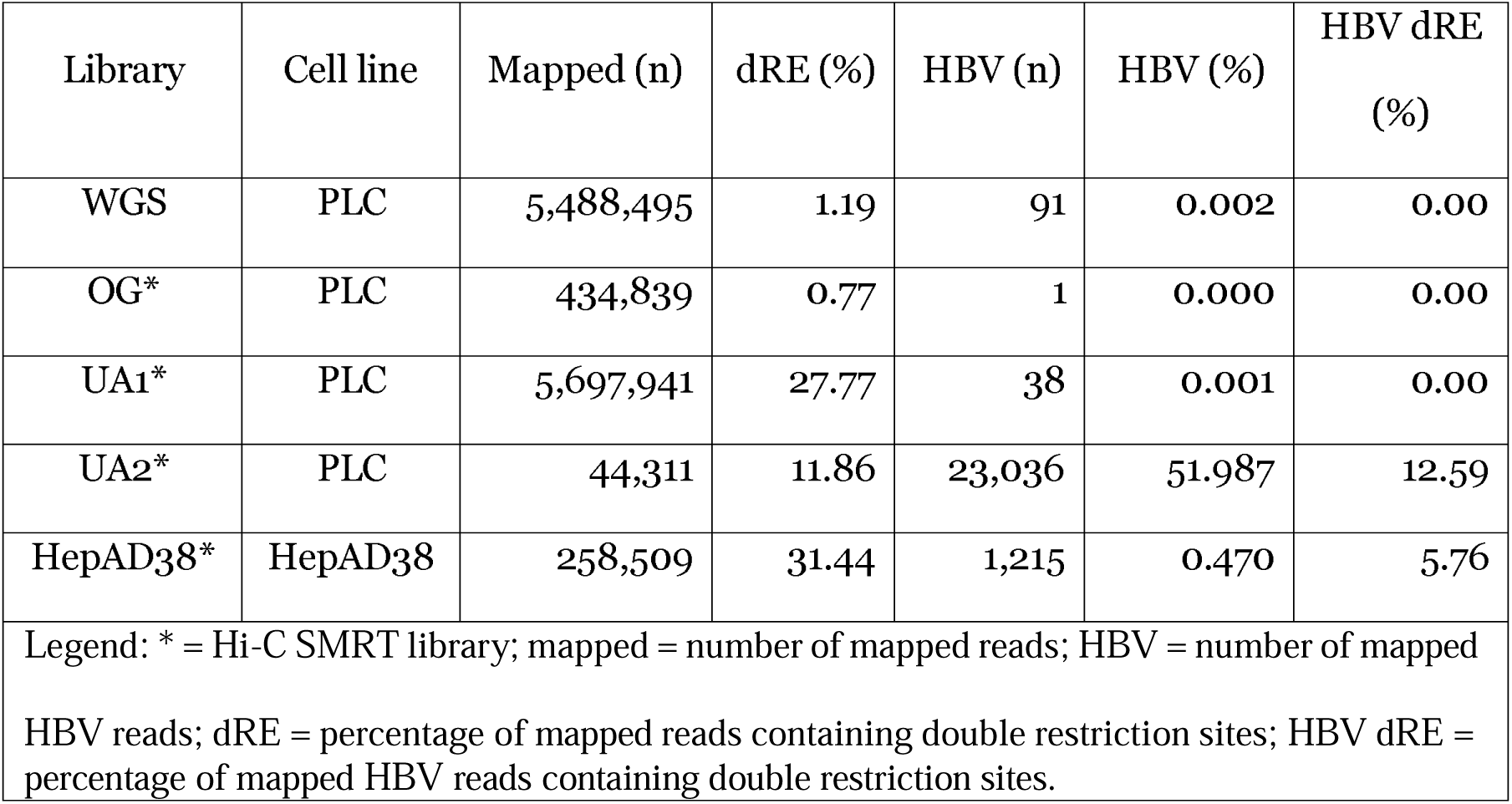
Read breakdown of each library.

| Library | Cell line | Mapped (n) | dRE (%) | HBV (n) | HBV (%) | HBV dRE (%) |
| --- | --- | --- | --- | --- | --- | --- |
| WGS | PLC | 5,488,495 | 1.19 | 91 | 0.002 | 0.00 |
| OG* | PLC | 434,839 | 0.77 | 1 | 0.000 | 0.00 |
| UA1* | PLC | 5,697,941 | 27.77 | 38 | 0.001 | 0.00 |
| UA2* | PLC | 44,311 | 11.86 | 23,036 | 51.987 | 12.59 |
| HepAD38* | HepAD38 | 258,509 | 31.44 | 1,215 | 0.470 | 5.76 |
| Legend: * = Hi-C SMRT library; mapped = number of mapped reads; HBV = number of mapped HBV reads; dRE = percentage of mapped reads containing double restriction sites; HBV dRE = percentage of mapped HBV reads containing double restriction sites. |  |  |  |  |  |  |

### HBV sequencing coverage within the Hi-C SMRT datasets

In addition to counting the number of reads, Integrative Genomics Viewer (IGV) was used to visualize the overall HBV genome coverage with the control and generated datasets: WGS, UA1, UA2, and HepAD38 (**Figure 5**) (Robinson et al. 2011; Thorvaldsdóttir et al. 2013; Robinson et al. 2017; Robinson et al. 2023). The coverage tracks over the HBV genome showed that the overall trend among the three datasets generated is similar, suggesting both Hi-C SMRT construction and HBV-target capture were not biased toward specific regions of the HBV genome. Note that there are multiple vertical colored lines across the coverage tracks suggesting nucleotide variation. This is not a concern since our datasets (UA1 and UA2) were high quality circular consensus sequencing (CCS) generated HiFi reads while the reads of the control WGS dataset were subreads which contain more sequencing errors. Furthermore, the reference sequence supplied for alignment was the HBV RefSeq NC_003977.2, HBV strain ayw, which is a genotype D (Okamoto et al. 1988). However, the HBV genotype present within the PLC cell line is genotype A (Ishii et al. 2020). The HBV coverage track of UA2, covering almost the entire HBV genome, suggested that our multi-genotype HBV capture probe panel successfully picked up all HBV sequences from the HBV genotype A. Encouragingly, the read coverage track of the HepAD38 cell line, containing the same HBV strain as the provided reference NC_003977.2, also showed that Hi-C SMRT construction and HBV-target capture probes worked well by covering the entire HBV genome (Ladner et al. 1997).

**Figure 5.**
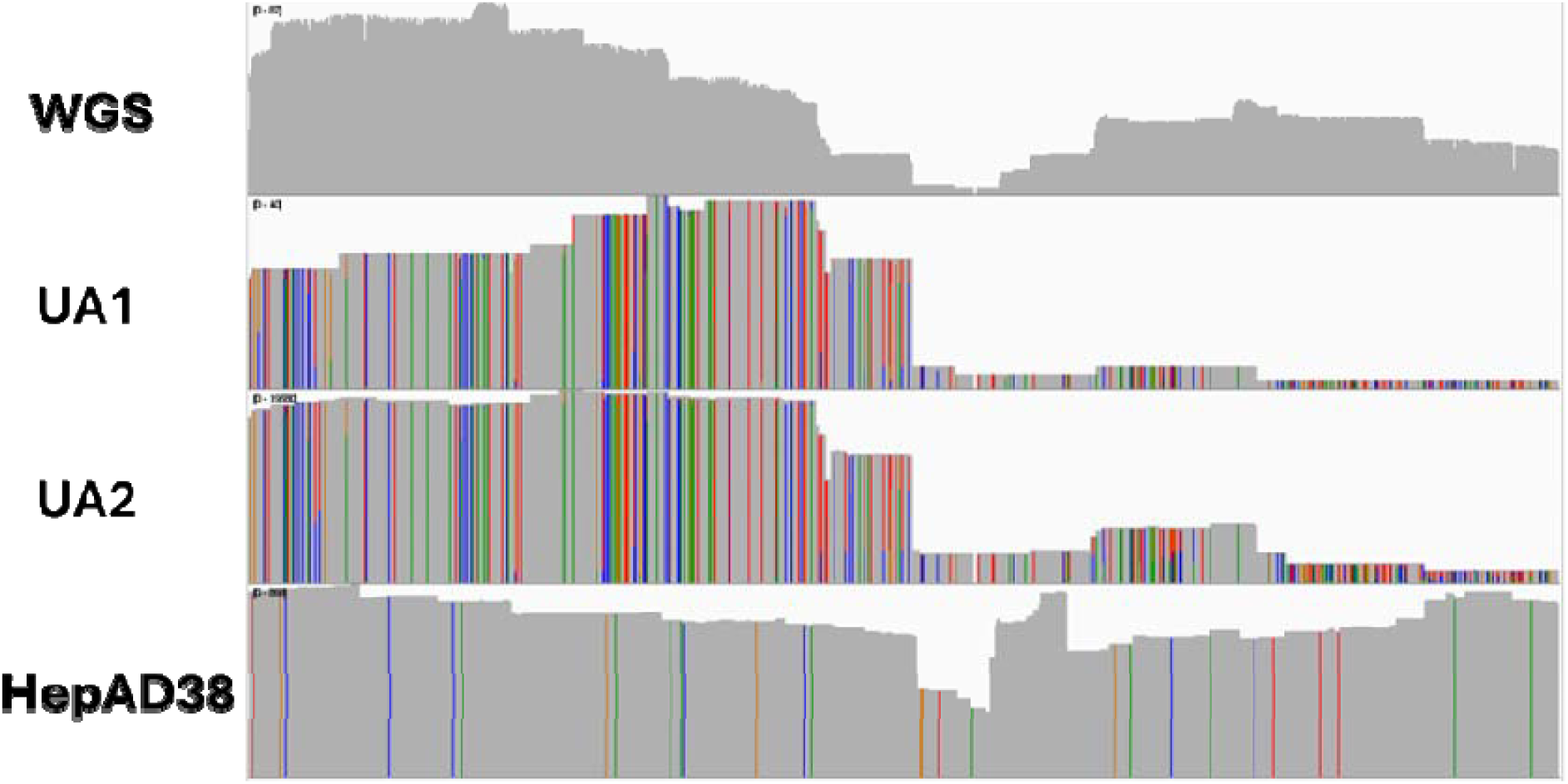
Overall sequencing coverage of the HBV reference genome, NC_003977.2 visualized using IGV. Vertical colored lines are default settings of the IGV, showing nucleotides that are different from the provided reference.

### Potential Hi-C contacts detected by Hi-C SMRT assay

Hi-C reads can be considered as artificially-generated chimeric reads with a specific connecting pattern, the double restriction site (dRE) sequence. Thus, to visualize HBV Hi-C reads, the HBV-containing reads were extracted and converted to browser extensible data (BED) files. The BED format captures and stores information of the alignment such as the coordinates of aligned read to the coordinates of the reference. Then, the reads were plotted along with the HBV genome using an R package, ggplot. Similar to IGV, each line represent a read that aligned to the reference genome (here, HBV, NC_003977.2) but color coded according to its chimeric partner on the human chromosome In other words, individual reads are plotted on the y-axis, the segment that mapped to HBV is presented as a track along x-axis according to the HBV genome region it covers, and colored coded based on which human chromosome the other segment aligned with. These plotted color-coded reads thus contain both chimeric Hi-C reads as well as HBV iDNA reads. The HBV reads from the reference WGS dataset (prepared without Hi-C) were plotted to serve as control to see if the same HBV iDNA sites were identified and if different sites were detected (**Figure 6**). From the plot, there were 9 different chimeric partners identified, suggesting there are at least 9 different HBV iDNAs presented in PLC cell line we were using. Next, we plotted HBV reads from UA1, which is a Hi-C SMRT library and identified 7 chimeric partners (**Figure 7**). The missing chimeric partners between WGS and UA1 could be coming from the low HBV sensitivity of the Hi-C SMRT assay and the additional chimeric partners, chr.5, could suggest Hi-C contacts. The plot of UA2, a Hi-C SMRT HBV-target captured library, revealed even more potential Hi-C contacts (**Figure 8**). More importantly, there were many dRE sites identified in these HBV-mapped reads from UA2, demonstrating a large number of Hi-C contacts. The high number of chimeric partners identified in HepAD38 could suggest pre-existing iDNA or Hi-C connections due to the *Hind*III site on the ptetHBV plasmid (**Figure 9**). To further focus on Hi-C reads, I filtered HBV-mapped reads with dRE and generated BED files of the filtered dRE-containing and HBV-containing reads (**Figures 10 and 11**) and found that after dRE filtering, there were overlapping chromosome partners between dRE filtered UA2 and WGS as expected (**Table 4**). Thus, we have demonstrated that our Hi-C SMRT HBV-capture assay is capable of picking up sequences containing all three target components including: dRE sequences, integrated HBV sequences; and discontinuous host sequences, indicative of topological contact.

**Figure 6.**
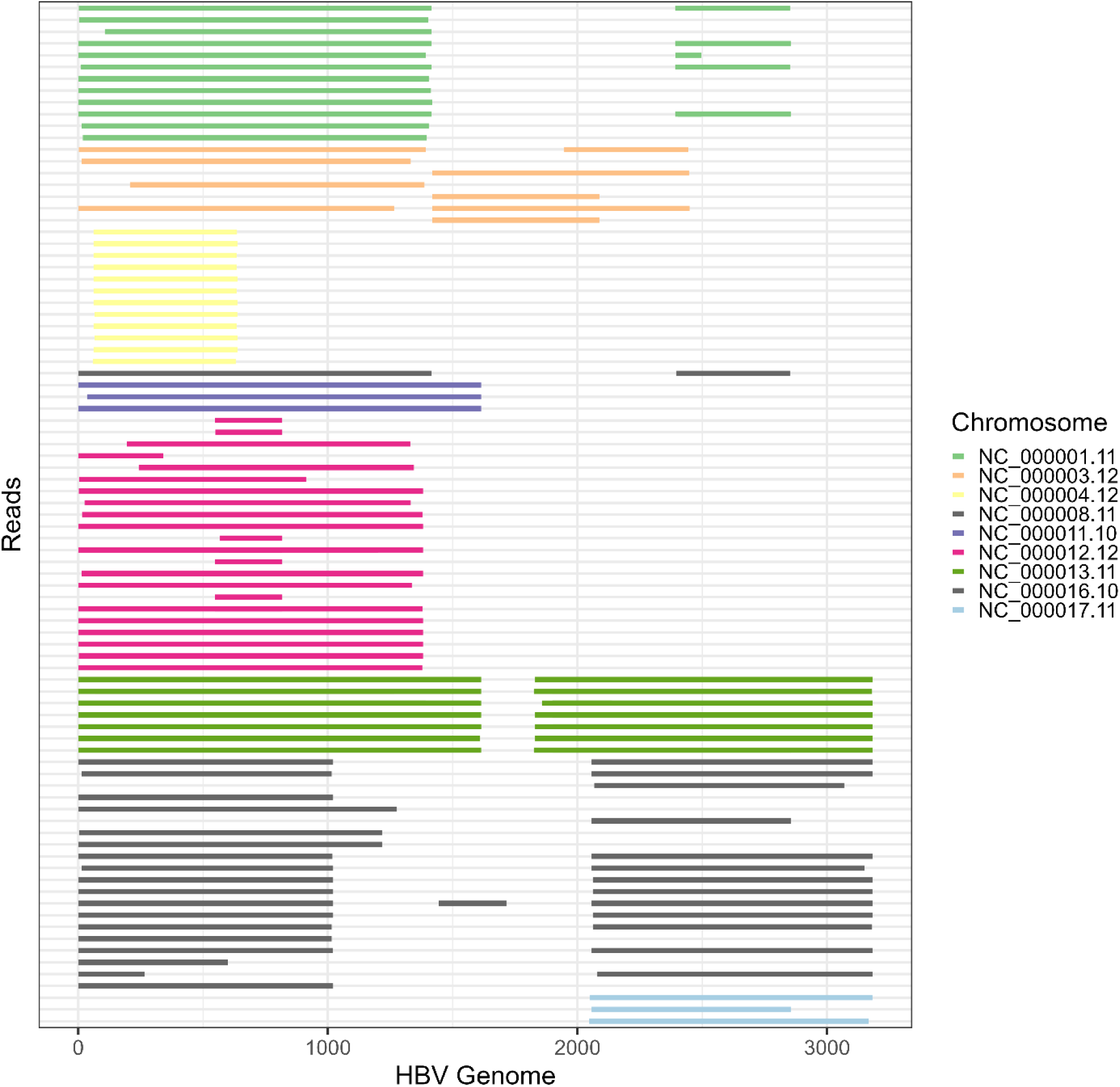
WGS chimeric read coverage across the HBV genome color coded with their chimeric chromosome partners. Y-axis represents individual reads and the x-axis represents the HBV genome. Individual reads are colored coded based on their chimeric partner on the human chromosome.

**Figure 7.**
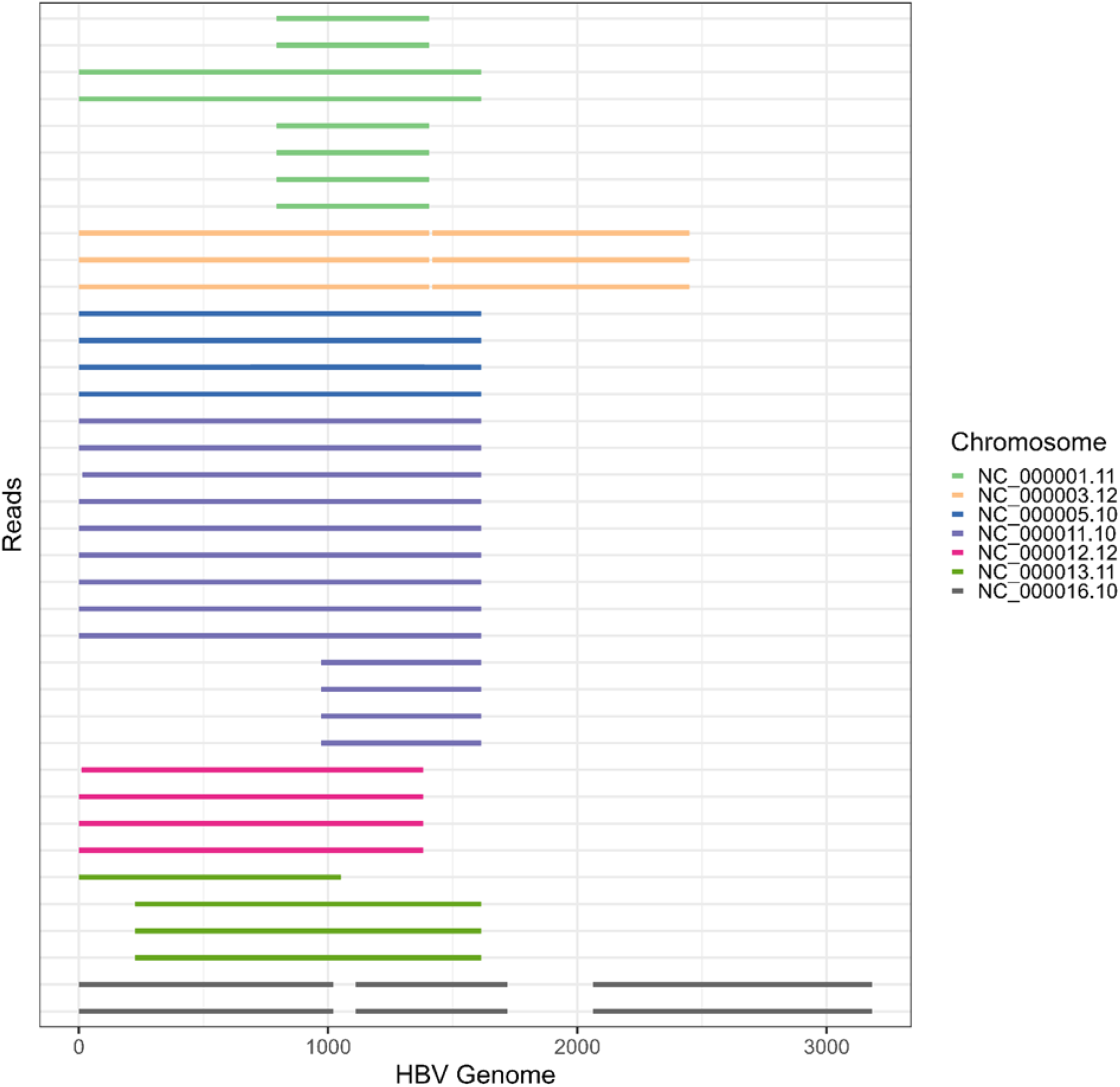
UA1 chimeric read coverage across the HBV genome color coded with their chimeric partners. Y-axi represents individual reads and the x-axis represents the HBV genome. Individual reads are colored coded bas on their chimeric partner on the human chromosome.

**Figure 8.**
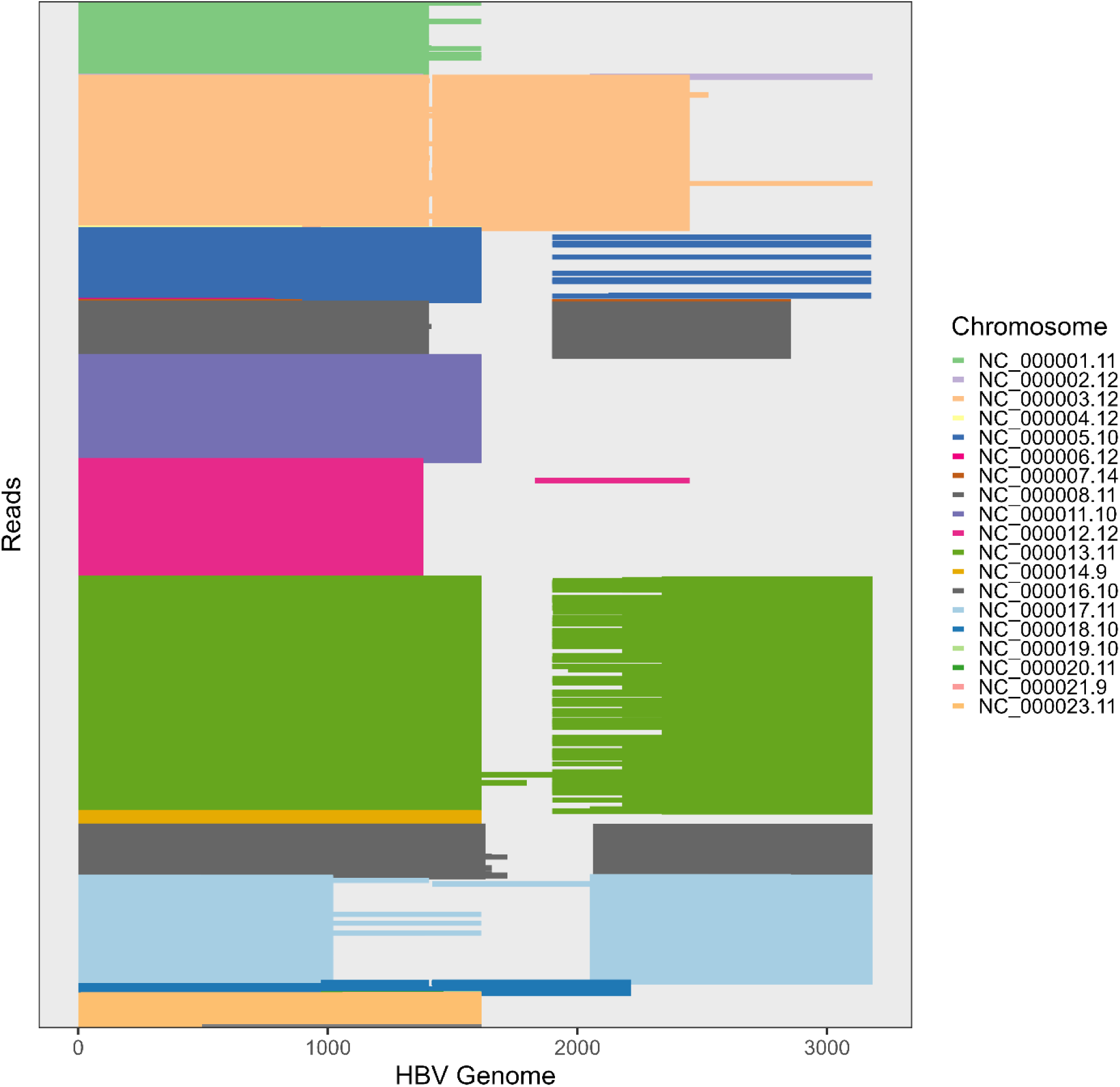
UA2 chimeric read coverage across the HBV genome color coded with their chimeric partners. Y-axi represents individual reads and the x-axis represents the HBV genome. Individual reads are colored coded bas on their chimeric partner on the human chromosome.

**Figure 9.**
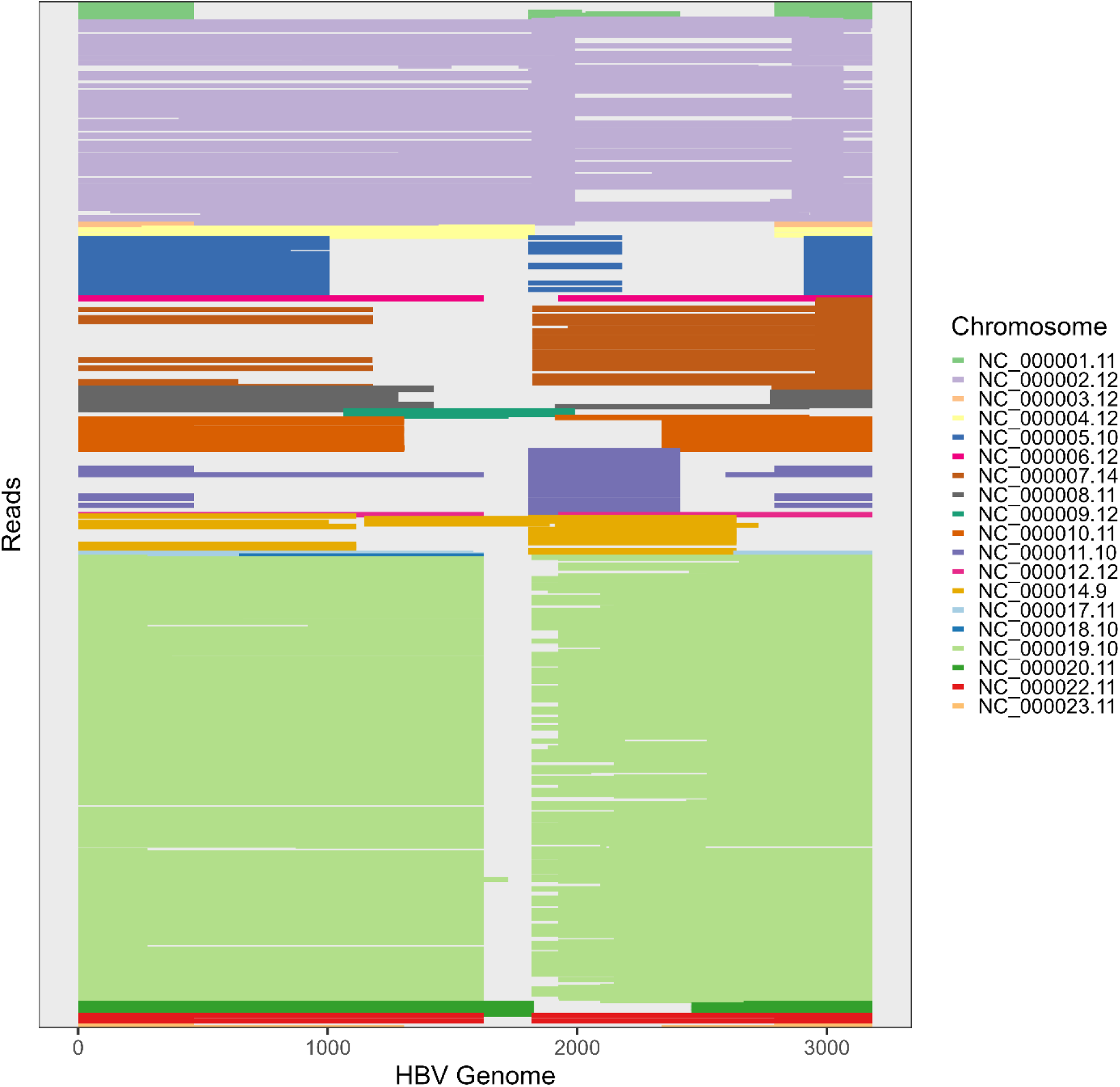
HepAD38 chimeric read coverage across the HBV genome color coded with their chimeric partners. Y-axis represents individual reads and the x-axis represents the HBV genome. Individual reads are colored coded based on their chimeric partner on the human chromosome.

**Figure 10.**
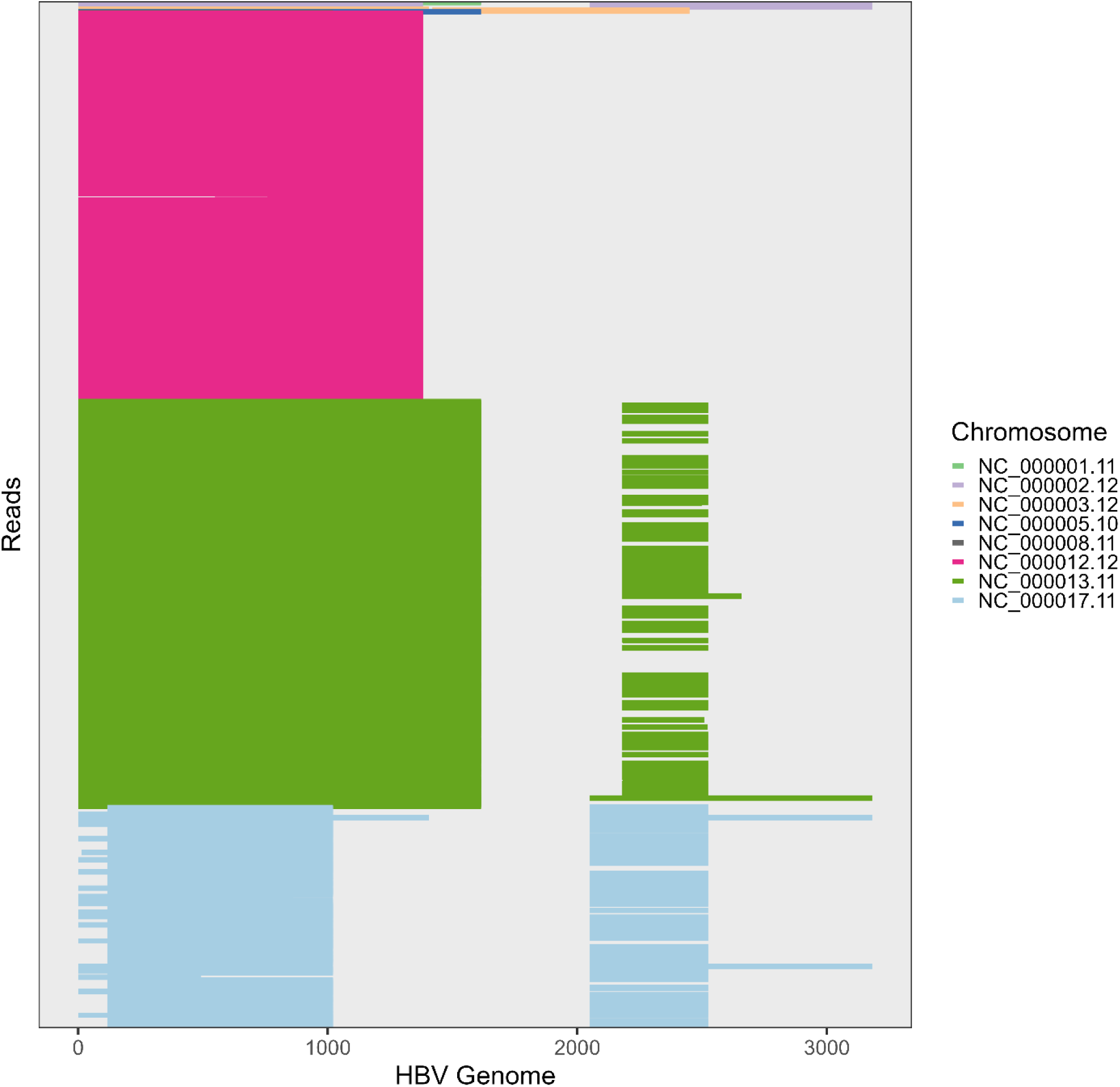
dRE filtered UA2 chimeric read coverage across the HBV genome color coded with their chimeric partners. Y-axis represents individual reads and the x-axis represents the HBV genome. Individual reads are colored coded based on their chimeric partner on the human chromosome.

**Figure 11.**
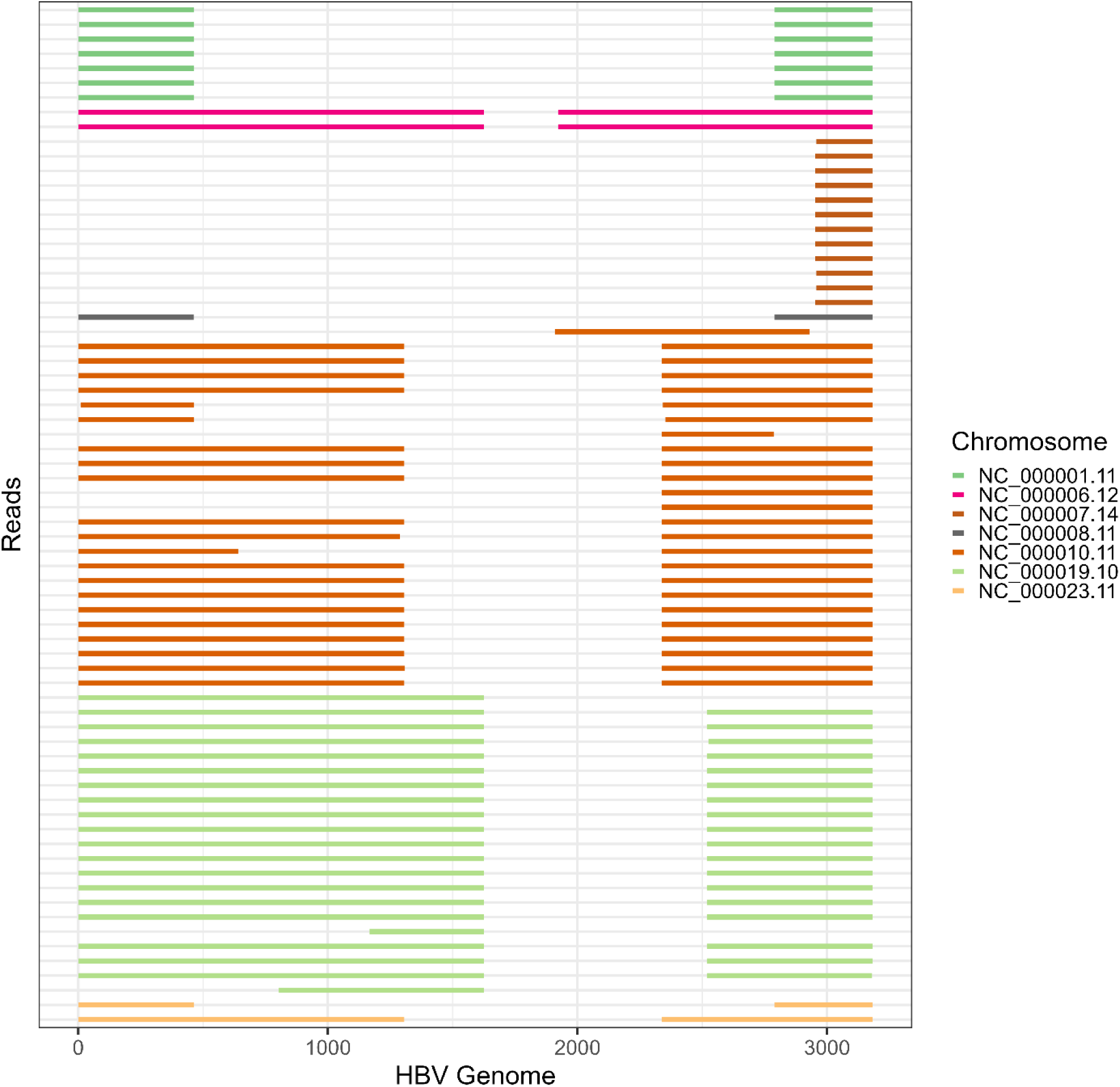
dRE filtered HepAD38 chimeric read coverage across the HBV genome color coded with their chimeric partners. Y-axis represents individual reads and the x-axis represents the HBV genome. Individual reads are colored coded based on their chimeric partner on the human chromosome.

**Table 4.**
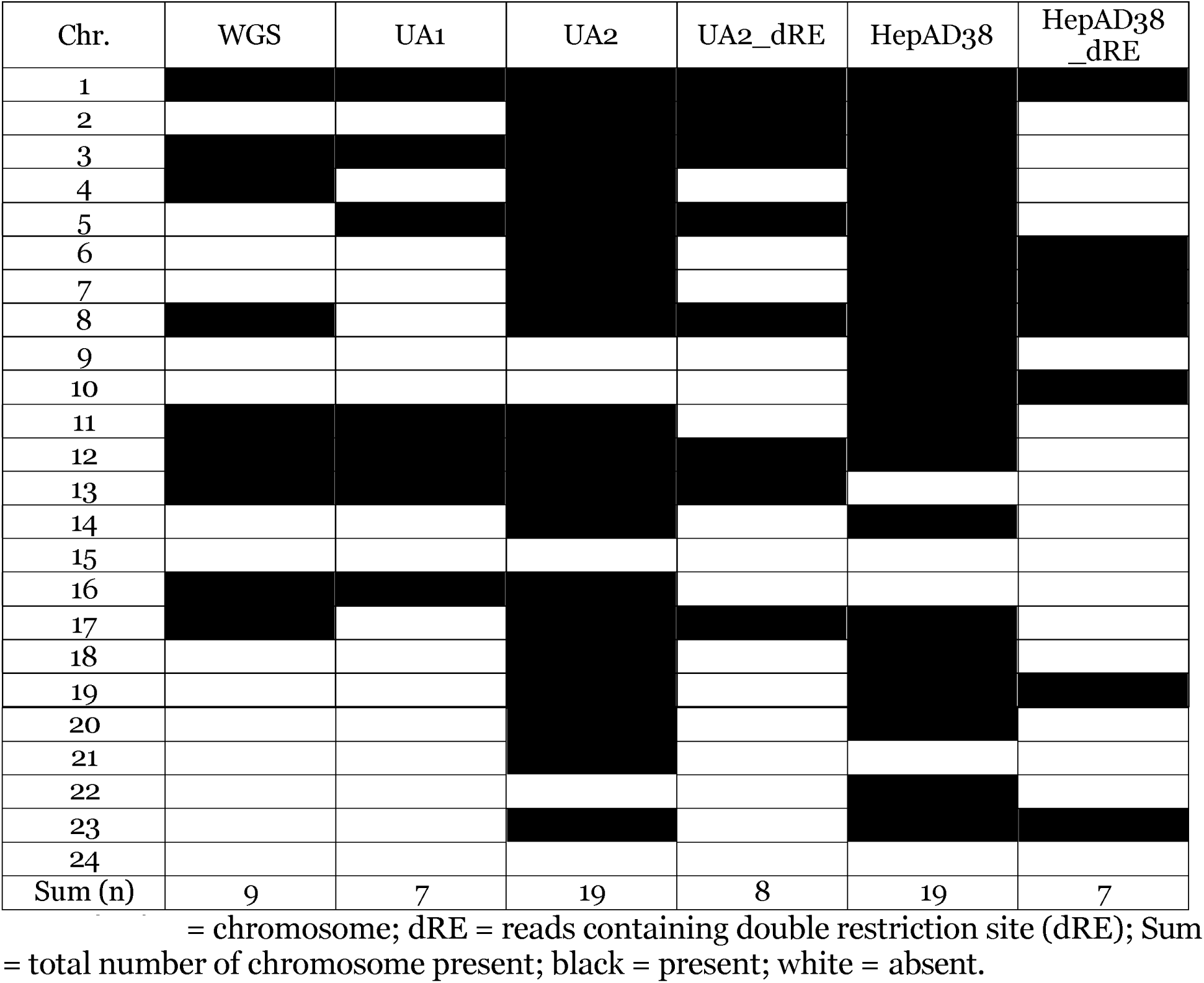
Summary of HBV chimeric reads’ chimeric partners.

### Visualization of the SMRT Hi-C reads

Although there are tools to visualize alignment results, such as IGV, these tools visualize reads using a reference sequence as backbone which is useful for viewing sequencing coverage of the reference (Fi**gure 12A**). However, reference-based position visualization is less straightforward in presenting the structure of the reads compared to query/read-based position visualization (**Figure 12B**). Thus, we converted the information from BAM and BED files (reference-based position) into query/read-based position information. With these data transformations, we were able to visualize the structure of each read, making it more straightforward to observe if individual dRE sequences were located between two non-genomically contiguous regions, thus indicating that the visualized read was the result of a Hi-C contact cut and ligation. Detailed scripts for these conversions are included in the Supplementary File under Section Read structure visualization.

**Figure 12.**
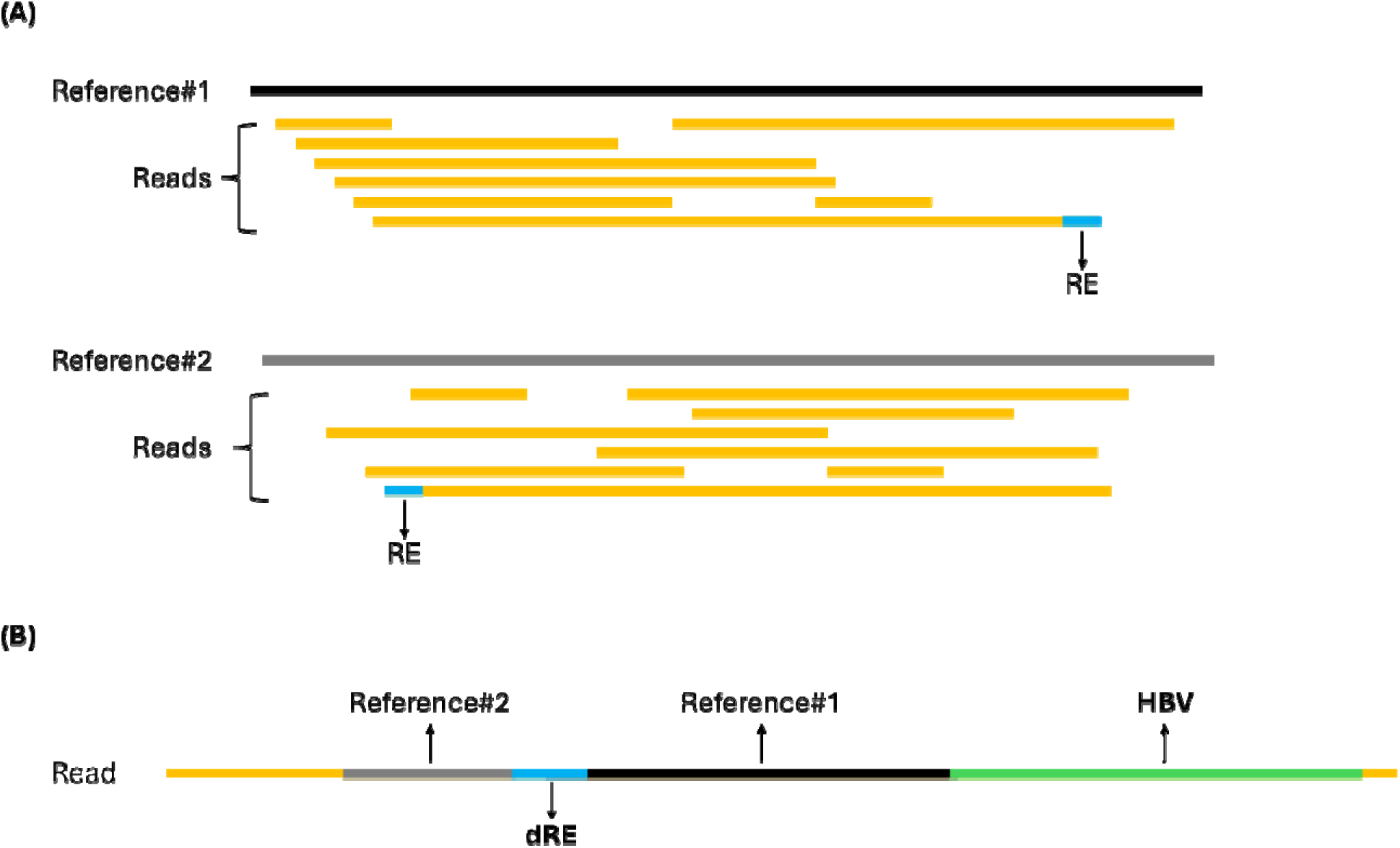
Alignment visualization strategies. (A) Alignment visualization using reference-based position information with tools such as IGV and Tablet. (B) Concepts of visualizing read structures using query/read-based position information. RE = restriction site; dRE = double restriction site.

## DISCUSSION

We have successfully incorporated multiple modifications into standard Hi-C procedures resulting in a novel long-read sequencing approach for studying 3D genomic interactions associated with integrated DNAs (**Figure 13**). By substituting rare cutter restriction endonucleases and integrating non-shearing probe hybridization capture methods designed for large templates, we successfully constructed and captured Hi-C libraries with average insert lengths of 5-6 Kb. Our post-sequencing analyses also indicated our capture probe panel effectively increased assay specificity and sensitivity towards the HBV genome without bias across its entire genome. The analytic results of our Hi-C SMRT libraries demonstrated their ability to detect and characterize Hi-C contacts resulting in chimeric reads containing dREs.

**Figure 13.**
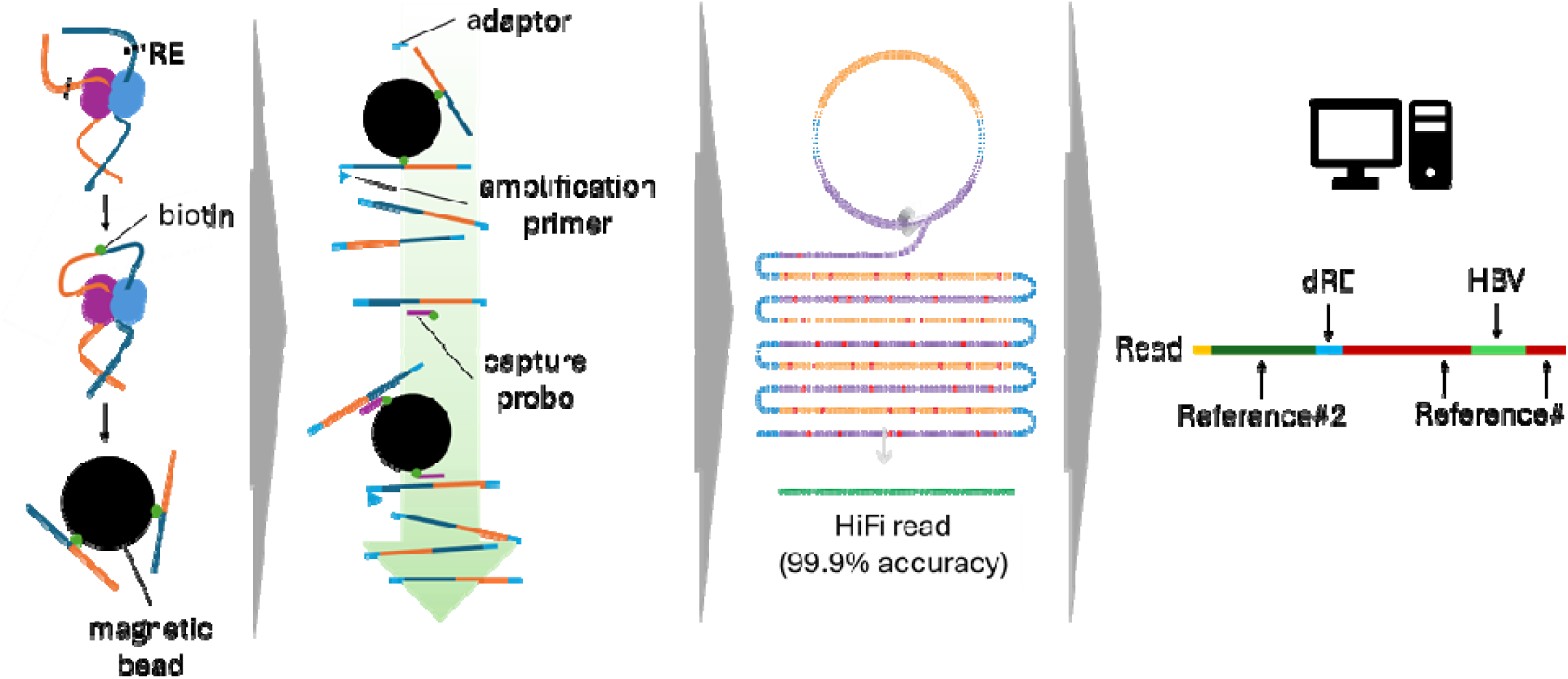
Hi-C SMRT workflow. Hi-C construction was performed with a customized restriction enzyme (RE) to ensure long DNA fragment length and intact target iDNA. Next, adaptor ligation amplification was performed to obtain the captured bead-bound Hi-C libraries followed by another long-DNA friendly hybridization pull-down step targeting HBV-containing sequences. The resultant Hi-C libraries are then subjected to SMRT library construction and sequencing. Finally, after sequence alignment and filtering, the structure of the potential Hi-C reads are visualized. dRE: duplicated *Hind*III restriction sites.

Additional layers of enrichment could be added to pursue higher sensitivity and specificity, however, the risk of obtaining over-amplified libraries should also take into considerations. Our processes involved two layers of amplifications, first to generate an off-capture-bead library and second to capture the targeted Hi-C library, although both are necessary, we did find seemingly duplicated reads in our data sets. Further optimization of the amplification steps or labeling the PCR-clonally expanded products by incorporating dual indexes during the process could be helpful. To further improve HBV-specific capture efficiency, a more intensive evaluation of each capture probe could be performed by determining their level of homology to the human genome and then re-evaluating the overall coverage after removing any highly human-homologous probes. We are aware that the long-read sequencing platforms provide lower coverage compared to short-read platforms and do not meet the suggested coverage for short-read high-confidence Hi-C contact report. However, long-read sequencing platforms, particularly the PacBio HiFi CCS sequencing technology, provide much greater and improved information including, fidelity, sequentially and spatially. With such high-quality sequencing results and optimized target pull-down approaches, we anticipate the Hi-C SMRT assay with read-lengths >20X that of short-read platforms would need, at most, 5% of the coverage required for standard short-read approach to confidently report Hi-C contacts. Although further evaluations should be performed to validate the contacts detected by our Hi-C SMRT assay, we anticipate the final established protocol will have the ability to report genomic contacts with high confidence levels based on the high-quality long-read sequencing data with much lower coverage levels required.

## METHODS

### Cell cultures

PLC/PRF/5 (PLC) cells were a generous gift from Dr. Tianlun Zhou (The Baruch S. Blumberg Institute, Doylestown, PA). PLC cells were cultured in Dulbecco’s Modified Eagle’s Medium (DMEM, Corning, Corning, NY) supplemented with 1X penicillin and streptomycin (VWR, Radnor, PA) and 10 % fetal-bovine serum (FBS, Gibco, Billings, MT) and incubated at 37 °C and 5 % CO2. HepAD38 was acquired from Dr. Haitao Guo’s Lab (University of Pittsburgh, Pittsburgh, PA). HepAD38 cells were cultured in DMEM/F12 (Gibco, Billings, MT) supplemented with 1X penicillin and streptomycin, 10% FBS, and 0.3 µg/mL of tetracycline hydrochloride (Research Products International, Mount Prospect, IL). Routine maintenance of the cell lines included refreshing culture medium every 3 to 4 days and passage of the cells when the culture reached a confluency of 80% to 90%, according to the recommendations of the American Type Culture Collection (ATCC, Manassas, VA). Detailed steps of cell passaging were as follows: (1) culture media was removed from the flask; (2) the culture flask (Falcon tissue culture treated flask, Corning, Corning, NY) was rinsed the with Dulbecco’s Phosphate Buffered Saline (DPBS, Gibco, Billings, MT); (3) trypsin (Gibco, Billings, MT) was added to form a thin layer covering the surface of the culture flask and incubated at 37 °C for 2 minutes; (4) fresh culture medium was added to neutralize trypsin; (5) the cell mixture was transferred to a new 15 mL conical tube and centrifuged at 1000 rpm for 1 minute; (6) the supernatant was discarded; (7) the cell pellet was resuspended in fresh medium and seeded into new flasks in a 1:3 to 1:6 ratio. Vials of cells were cryopreserved at approximately 1 million cells per vial in 5 % dimethyl sulfoxide (DMSO) supplemented DMEM.

### Hi-C library construction

Freshly harvested or frozen cell pellets of PLC cells and HepAD38 cells were used for Hi-C single molecule, real-time (SMRT) library construction. Hi-C construction was performed using Proximo Hi-C (human) Kit (Phase Genomics, Seattle, WA) with modifications with the choice of restriction enzyme used for DNA digestion. *Hind*III-HF was purchased from New England BioLab (NEB, Ipswich, MA) to replace restriction enzyme in the standard Hi-C protocol. An R package, DECIPHER, was used for *in silico* estimates of restriction enzyme digestions. The human reference genome (GRCh38, hg38) and HBV reference genome (NC_003977.2) were downloaded from National Center for Biotechnology Information (NCBI) and used for *in silico* analyses.

### Adaptor ligation and amplification of the Hi-C library

Adaptor ligation and amplification steps were adapted from PacBio’s (Menlo Park, CA) “Multiplex Genomic DNA Target Capture Using IDT xGen Lockdown Probes” to replace the steps of on-bead library preparation described in the Proximo Hi-C (human) kit. Supplies adapted from PacBio protocol include KAPA Hyper Prep Kits (Roche, Indianapolis, IN), Dynabeads M-270 Streptavidin (Invitrogen, Waltham, MA), forward universal adaptor oligo, reverse universal adaptor oligo, PacBio universal primer, and TaKaRa LA Taq DNA Polymerase Hot-Start Version (TaKaRa, San Jose, CA). The forward, reverse, and universal primers (Supplementary Table 2) were purchased from IDT (Newark, NJ).

### HBV hybridization capture

Twist Bioscience’s (South San Francisco, CA) customized hybridization capture kit was used for HBV-targeted DNA capture. Custom probes were designed by Twist Bioscience based on the 42 HBV genotypic references sequences published by McNaughton et al., 2020.

### SMRT library construction

The PacBio HiFi express template preparation kits 2.0 and 3.0 (PacBio, Menlo Park, CA) were used for SMRT library construction.

### Sequencing and analysis

Pre-sequencing quality controls were performed using the Qubit dsDNA HS assay (Invitrogen, Waltham, MA), the TapeStation Genomic DNA ScreenTape assay (Agilent, Santa Clara, CA), and an Agilent Femto Pulse (Invitrogen, Waltham, MA) for library concentrations and size distributions. DNA sequencing runs were performed on a PacBio Sequel IIe with a movie time of 30 hours. FastQC was used for basic assessments of sequencing quality metrics. PacBio’s SMRT link (PacBio, Menlo Park, CA) was used for obtaining the sequencing run quality. The Human reference genome (GRCh38, hg38) and the HBV reference genome (NC_003977.2) were downloaded from National Center for Biotechnology Information (NCBI) and used for sequence analyses. Minimap2 and samtools were used for alignment (Li 2018; Danecek et al. 2021; Li 2021). Linux utility functions, R, and RStudio were used for data analysis and visualization. The interactive genomics viewer (IGV) was used to view the sequencing coverage. Tools utilized on our Linux platform for sequence processing included: FastQC; minimap2; samtools; bedtools; seqkit; seqtk; and GATK (Quinlan and Hall 2010; Li 2012; Shen et al. 2016; Li 2018; Van der Auwera and O’Connor 2020; Danecek et al. 2021; Li 2021). Tools and packages utilized in R included: pacman, CRAN, cli, devtools, tidyverse, ggraph, igraph, RColorBrewer, tidyr, knitr, ggpmisc, gggenes, BiocManager, and DECIPHER.

### Referencing whole genome sequencing data set

The whole genome sequencing (WGS) data of PLC cell line was downloaded from NCBI Sequence Read Archive (SRA; Run: SRR14087064; BioProject: PRJNA717995).

## DATA ACCESS

The demultiplexed sequencing data that support the establishment of this study have been submitted to the NCBI BioProject database (https://www.ncbi.nlm.nih.gov/bioproject/) under accession number PRJNA# (in prep.).

## ACKNOWLEDGEMENTS

This work was supported by the Oskar Fischer Project, a gift from the James Truchard Philanthropies; JBS Science Inc; and NIH grants DC02148 and DK082316 to GDE. Yih-Ping Su was the recipient of a pre-doctoral fellowship supported by JBS Science Inc. Dr. Tianlun Zhou gifted us PLC/PRF/5 cells. Dr. Haitao Guo gifted us HepAD38 cells. We wish to thank Joshua Chang Mell for fruitful discussions.

## AUTHOR CONTRIBUTIONS

Yih-Ping Su designed and performed the bulk of the experiments; developed the informatic pipeline, analyzed and curated the data including preparing the graphic visualizations, and prepared original draft. Garth D. Ehrlich conceptualized the project and edited the entire manuscript with the discrete laboratory and computational work performed by Yih-Ping Su and Joshua P. Earl. Joshua P. Earl contributed to Yih-Ping Su’s bioinformatic pipeline building supervision and training. Azad Ahmed and Bhaswati Sen contributed to sequencing set-up and related optimization. Joshua P. Earl and Samuel Czerski performed post-sequencing sample demultiplex processes and sequencing run quality report. Garth D. Ehrlich acquired the funding.

